# Kinetic proofreading as a mechanism for transcriptional specificity in living human cells

**DOI:** 10.64898/2026.03.17.711757

**Authors:** Jee Min Kim, David A. Ball, Thomas A. Johnson, Chris D’Inzeo, Hyeon Jin Cho, Laurent Ozbun, Tatiana S. Karpova, Gianluca Pegoraro, Daniel R. Larson

## Abstract

How target genes selectively respond to their specific transcription factors (TF) amid the vast excess of non-specific TFs in the nucleus remains a fundamental question in gene regulation. Here, we develop an integrated single-molecule imaging framework to quantitatively link TF dynamics with nascent transcription kinetics at endogenous gene loci to address the transcription factor specificity. Using endogenously Halo-tagged glucocorticoid receptor (GR), we show that ligand activation rapidly increases GR chromatin binding and residence times without substantially altering search kinetics. Live-cell nascent RNA imaging reveals that GR is essential for Dex induction of *ERRFI1* transcription via increased burst frequency. Gene locus-specific, dual-color tracking demonstrates that GR display longer residence times in general near its target gene *ERRFI1* compared to the non-target *MYH9* locus, consistent with a kinetic proofreading model. A high-throughput imaging-based CRISPR screen identifies ATP-dependent processes, including neddylation and chromatin remodeling, and similarly, acute inhibition of the TFIIH XPB selectively impairs *ERRFI1* transcription while sparing *MYH9*, implicating ATPase activity in GR target discrimination. In all, these findings establish that promoters function as dwell-time rather than occupancy detectors, discriminating specific from non-specific TF interactions through energy-dependent mechanisms.

## Introduction

Transcription factors are key regulators of gene expression that define a cell’s identity, function, and response to external cues. Single-molecule studies over the past decades have uncovered how transcription factors in living cells interact with chromatin in a highly dynamic manner (Chen et al., 2014; Gebhardt et al., 2013; Mazza et al., 2012; Swinstead et al., 2016). Concurrently, transcription occurs in a series of episodic bursts of nascent mRNA synthesis interspersed with refractory periods of inactivity, a phenomenon that is conserved from yeast to mammals (Bothma et al., 2014; Raj & van Oudenaarden, 2008; Rodriguez & Larson, 2020). How dynamic transcription factor–chromatin interactions are quantitatively linked to stochastic transcriptional bursting to produce specific, robust gene expression profiles remains an ongoing question in the field.

Central to this long-standing question is the problem of transcriptional specificity: how target genes selectively respond to their specific transcription factors amid the vast excess of non-specific factors present in the nucleus. In yeast, recent work has begun to address this question by simultaneously visualizing transcription factor binding and transcriptional output at a single target gene, revealing the role of cooperative transcription factor dynamics in shaping bursting kinetics (Donovan et al., 2019; Pomp et al., 2024). While studies have investigated transcription factor kinetics in relation to transcriptional output in mammalian cells (Popp et al., 2021; Stavreva et al., 2019), much less is known about the kinetic signatures of endogenous transcription factors in mammalian cells that discriminate between target and non-target genes within their native gene landscape.

There has been considerable recent work examining different dynamic models of transcription (reviewed in (Kim & Larson, 2025)). However, to our knowledge there have been few if any measurements which directly address the question of specificity in living cells. Historically, transcription kinetics were described using two-state telegraph and three-state models, usually assuming thermodynamic equilibrium, thereby imposing an upper limit on specificity (Harper et al., 2011; Ko, 1991; Peccoud & Ycart, 1995; Tantale et al., 2016). Simply put, if the difference between specific and non-specific dwell times is ∼ 100-fold but the non-specific factors are 100-fold more abundant, the relative occupancy on *cis*-regulatory elements in chromatin by equilibrium models will be ∼ 1:1 (Herschlag & Brad Johnson, 1993). In contrast, the kinetic proofreading model, a non-equilibrium framework originally proposed by Hopfield and Ninio, can enhance specificity beyond this limit through irreversible, energy-consuming steps that discriminate between specific and non-specific factors (Biddle et al., 2019; Boeger, 2022; Hopfield, 1974; Ninio, 1975; Shelansky et al., 2024). Moreover, the molecular identity of the energy-dependent processes that might underpin transcriptional proofreading in mammalian cells remains largely unknown.

We examine the glucocorticoid receptor (GR; encoded by *NR3C1*), which offers an ideal system to address these questions. As a ligand-inducible transcription factor, GR translocates to the nucleus upon steroid binding and engages glucocorticoid response elements to modulate target gene transcription, enabling precise control of activation. Single-molecule tracking studies have revealed that GR–chromatin interactions are dynamic (McNally et al., 2000; Voss & Hager, 2014), with residence times following a power law distribution that reflects a broad continuum of binding affinities and mobility-dependent dwell times (Garcia et al., 2021; Stavreva et al., 2019; Wagh et al., 2023).

Here, we develop an integrated experimental framework to quantitatively link GR single-molecule dynamics with nascent transcription kinetics at endogenous target and non-target genes. We show that GR activation by the synthetic agonist dexamethasone increases burst frequency at *ERRFI1* without substantially altering burst duration, and that this transcriptional activation requires the physical presence of GR. Fast and slow tracking reveals that Dex induces rapid, global increases in GR chromatin binding and residence times without major changes in search kinetics. Locus-resolved tracking demonstrates that GR molecules proximal to the *ERRFI1* target gene exhibit longer residence times than those near the non-target *MYH9* locus, consistent with a kinetic proofreading model in which energy-dependent steps selectively stabilize productive transcription factor–chromatin interactions at target genes. To identify the molecular machinery underlying this specificity, we performed a high-throughput imaging-based CRISPR screen and discovered that *ERRFI1*-specific regulators include ATP-dependent processes, such as neddylation, chromatin remodeler, and Mediator. Together, our findings establish a mechanistic link between transcription factor dwell times and gene-specific transcriptional output and identify candidate molecular effectors of kinetic proofreading in inducible transcription for further study.

## Results

### Inducible GR activation defines *ERRFI1* as a target and *MYH9* as a non-target gene to dissect kinetic signatures of transcriptional specificity

To study inducible gene transcription in a native genomic context, we first focused on the nuclear receptor superfamily, specifically glucocorticoid receptor (GR), retinoid X receptor (RXR), retinoic acid receptor alpha (RARα), and peroxisome proliferator-activated receptor gamma (PPARγ) nuclear receptors. Members of this steroid-inducible transcription factor family are activated upon ligand binding and induce transcriptional responses through direct interaction with specific genomic loci (Fig 1A).

**Figure 1.**
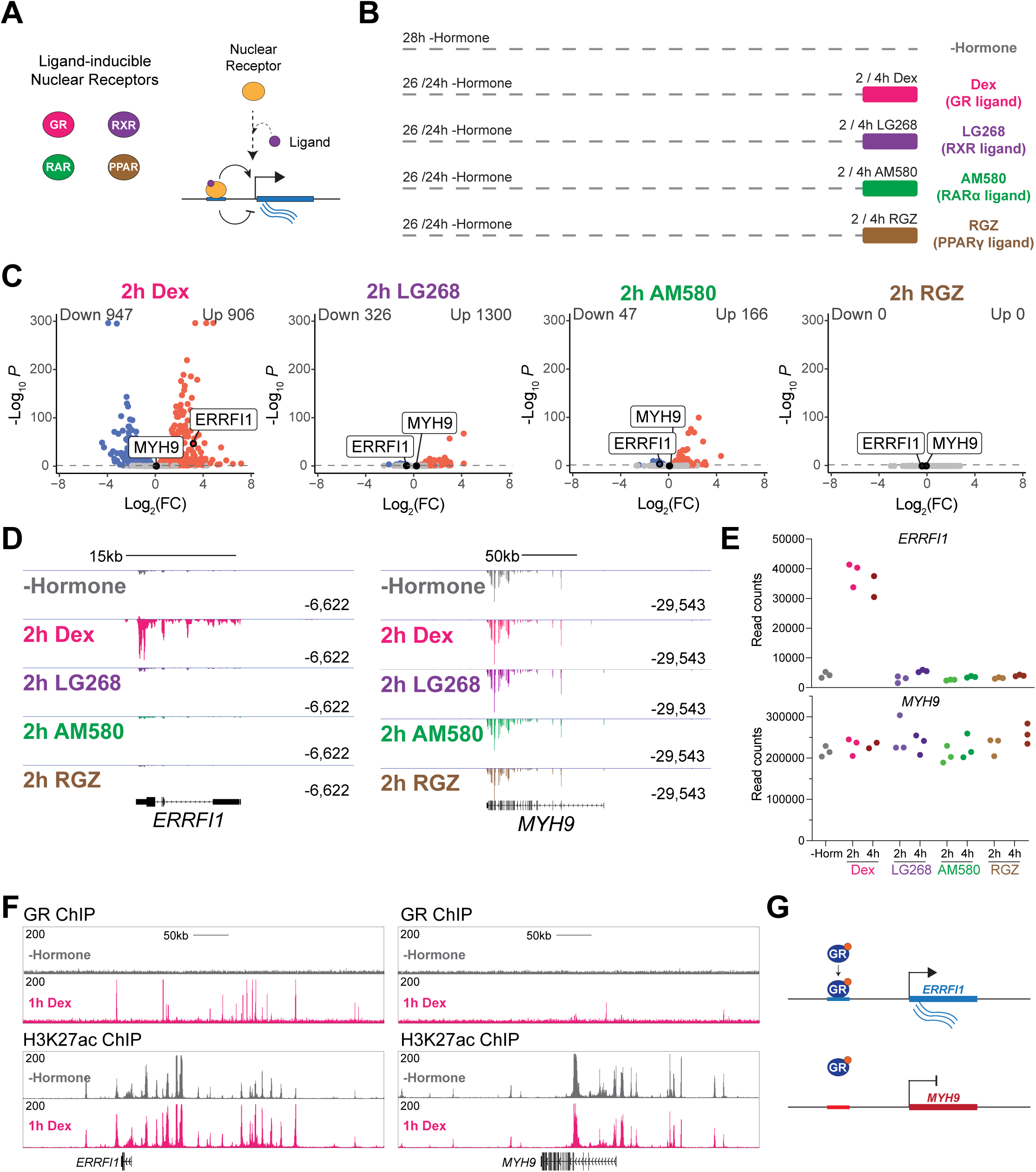
GR ligand-inducible transcription factor acts specifically to *ERRFI1* and not *MYH9* gene. (A) Schematic for transcriptional regulation by ligand-inducible nuclear receptors. (B) Experimental designs for either 2 or 4h activation of GR, RXR, RARa, and PPARy nuclear receptors using synthetic ligands in human bronchial epithelial cells. (C) Volcano plots showing differentially expressed genes by RNA-seq after activation of nuclear receptors 2h post-treatment (dashed lines indicate adj p-val threshold of 0.05). (D-E) RNA-seq genome browser tracks (D) and read count values (E) at the *ERRFI1* and *MYH9* loci at the indicated ligand treatments. (F) GR-ChIP and H3K27ac-ChIP genome browser tracks at *ERRFI1* in the absence and presence of Dex. (G) Summary of GR transcription factor acting as a specific transcription factor for *ERRFI1* and non-specific transcription factor for *MYH9* gene.

To look for functionality and target genes, we first tested the effects of activating GR, RXR, RARα, and PPARγ nuclear receptors in human bronchial epithelial cells, an immortalized and non-transformed human cell line, by treating them with synthetic ligands for 2 or 4 h (Fig 1B). To maximize target gene induction, cells were hormone-deprived for 24 h prior to ligand treatment. By RNA-seq, we found various extents of global transcriptional changes under GR, RXR, and RARα conditions, suggesting these transcription factors are active in HBECs as shown previously (Fig 1C). PPARγ ligand did not induce any transcriptional changes, suggesting that this transcription factor may not be active under our experimental conditions.

We decided to focus on two genes, *ERRFI1* and *MYH9*: *ERRFI1* is a GR-responsive gene that is strongly activated by GR ligand, Dexamethasone (Dex), whereas *MYH9* gene, despite its robust transcriptional activity in all conditions tested, is GR-insensitive, providing a stringent negative control within the same cellular milieu (Fig 1D, 1E, S1). Notably *ERRFI1* does not respond to other nuclear receptor ligands. Additionally, we performed ChIP-seq for GR and histone mark H3K27ac to show that GR occupies enhancer elements flanking the *ERRFI1* gene in a Dex-dependent manner, whereas no substantial GR occupancy is detected by Dex treatment for the *MYH9* gene despite having active enhancer elements proximal to the gene (Fig 1F). In all, these results provide evidence that GR acts as a specific transcription factor for the *ERRFI1* gene and a non-specific transcription factor for the *MYH9* gene (Fig 1G).

To simultaneously probe the kinetics of transcription factor and gene transcription at the endogenous level, we inserted HaloTag at the N-terminus of the GR gene locus (*NR3C1*) using CRISPR/Cas9 genome engineering. Endogenous tagging was performed in a cell line containing a 24x repeat of MS2 stem loop integrated in the first intron of the *ERRFI1* gene, where GR as expected displayed Dex-dependent nuclear translocalization (Fig 2A-B) (Wan et al., 2021). We also generated a cell line stably expressing Halo-GR construct by lentiviral transduction with a cell line containing 24X-MS2 integrated in the first intron of *MYH9* and the *ERRFI1*-MS2 cell line mentioned above (Fig S2).

**Figure 2.**
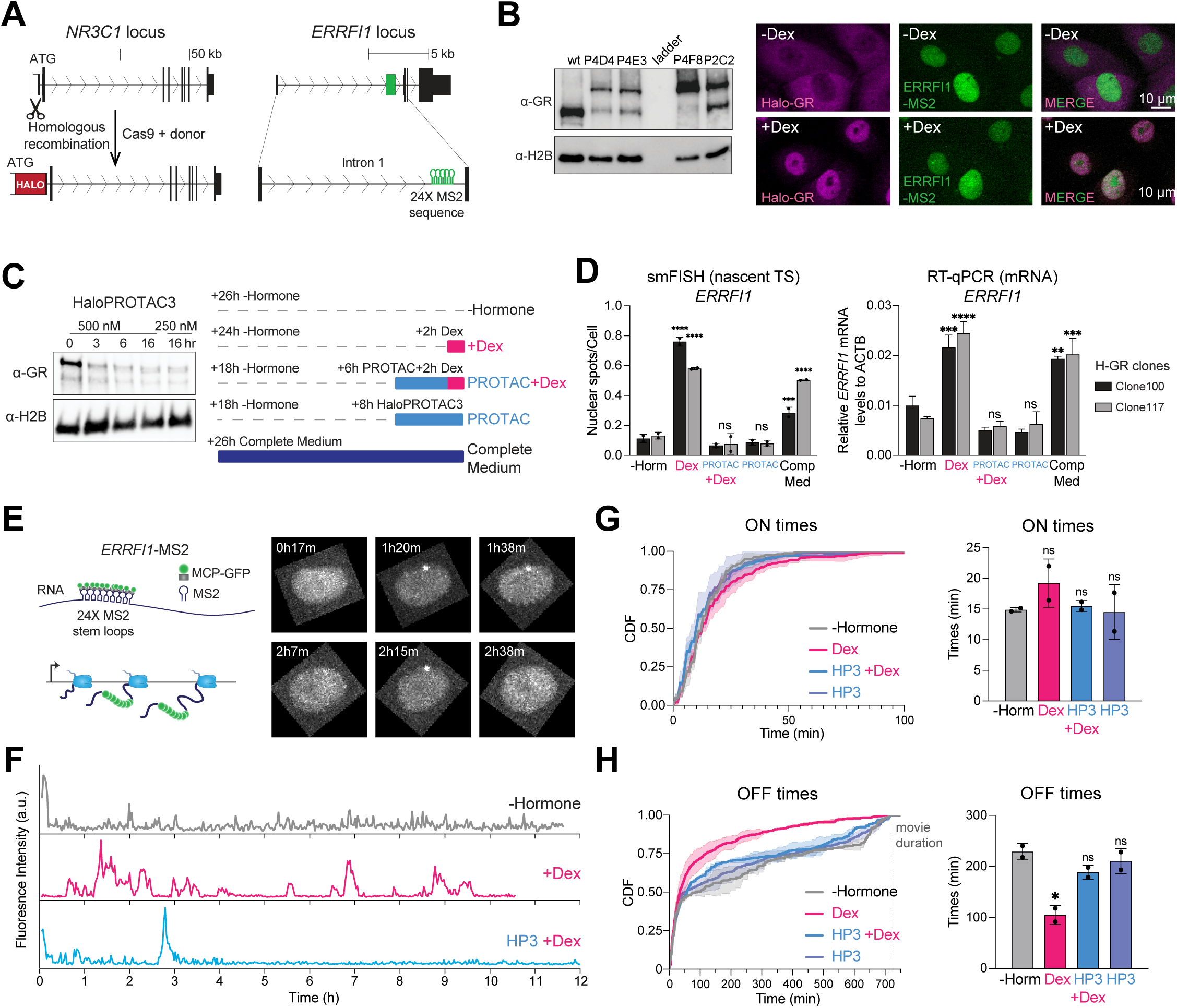
GR activates *ERRFI1* nascent transcription by decreasing OFF times by dex induction. (A) Schematic of endogenous HaloTagging of GR via CRISPR-Cas9 in a cell line containing 24×MS2 stem loops at the native *ERRFI1* locus (Wan et al., 2021). (B) Western blot analysis showing heterozygous and homozygous Halo-GR single clones (left) and confocal microscope images of Halo-GR cells in -Dex and +Dex conditions (right). (C) A time course showing degradation of Halo-GR using HaloPROTAC3 by Western blot (left). Experimental design to test the effects of GR degradation on *ERRFI1* gene induction by Dex (right). (D) Quantification of transcriptionally active *ERRFI1* alleles (left) and ERRFI1 mRNA levels (right) by smFISH and RT-qPCR, respectively. (E) Schematic of nascent RNA imaging using MS2 system (left), and representative time-lapse images of a nuclei at different timepoints (right). (F) Example time traces of nascent transcription imaging of the *ERRFI1* gene. (G-H) CDF plots (left) and quantification of each experimental condition (right) for *ERRFI1* ON (G) and OFF (H) times.

To validate specificity of GR on *ERRFI1* gene transcription at the nascent and mature RNA level, we used HaloPROTAC3, a small compound that covalently binds to HaloTag and recruits E3 ligase to degrade Halo-fusion proteins, in our homozygously tagged Halo-GR cell line (Buckley et al., 2015). We found substantial and saturating levels of GR degradation by 6 hr of HaloPROTAC3 treatment (Fig 2C). Using smFISH and RT-qPCR to detect nascent transcription site and total mRNA levels, respectively, we found rapid degradation of GR abrogates Dex induction of *ERRFI1* in two independent homozygous Halo-GR cell lines, establishing GR’s necessity for *ERRFI1* transcriptional activation (Fig 2D).

Live-cell, nascent RNA imaging of MS2/MCP-GFP was next performed to directly observe *ERRFI1* transcriptional bursting kinetics (Fig 2E). Using fluorescence intensity traces over time, we captured transcriptional ON and OFF times using Hidden Markov Model (HMM)-detection of ON peaks (Fig 2F). Whereas Dex-treated cells exhibit significantly shorter OFF times compared to hormone-starved cells, hormone-starved and HaloPROTAC3-treated cells exhibit similar transcriptional bursting kinetics of ON and OFF times (Fig 2H). Minimal changes were observed for ON times, suggesting that Dex primarily increases *ERRFI1* burst frequency rather than burst duration (Fig 2G), as has been observed previously for steroid receptors (Rodriguez et al., 2019). In total, these results support the necessary role GR plays in the nascent transcription activation of *ERRFI1* upon Dex induction.

### Global GR single-molecule dynamics reveal selective modulation of off-rates upon ligand activation

Having established a system in which transcription factor–gene target and non-target pairs can be directly compared, we next examined how GR binding kinetics globally change under inactive (-Dex) and active (+Dex) states. By capturing 10 ms-frame rate movies under continuous laser illumination (“fast tracking”), we directly measured biophysical parameters of single-molecule diffusivity (Fig 3A). Endogenous tagging enabled these measurements to reflect steady-state chromatin-binding frequency of the GR population. Single-molecule trajectories revealed heterogeneous GR behaviors within individual nuclei (Fig. 3B), with ligand activation inducing a marked increase in the chromatin-bound fraction from 4.3% to 32.7% (Fig. 3C). Time-course measurements at 15-min intervals showed that GR binding reaches steady state within 30 min of Dex addition, which correlates with activation of the GR target *ERRFI1* gene during the first 3 h of Dex induction (Fig 3D-F).

**Figure 3.**
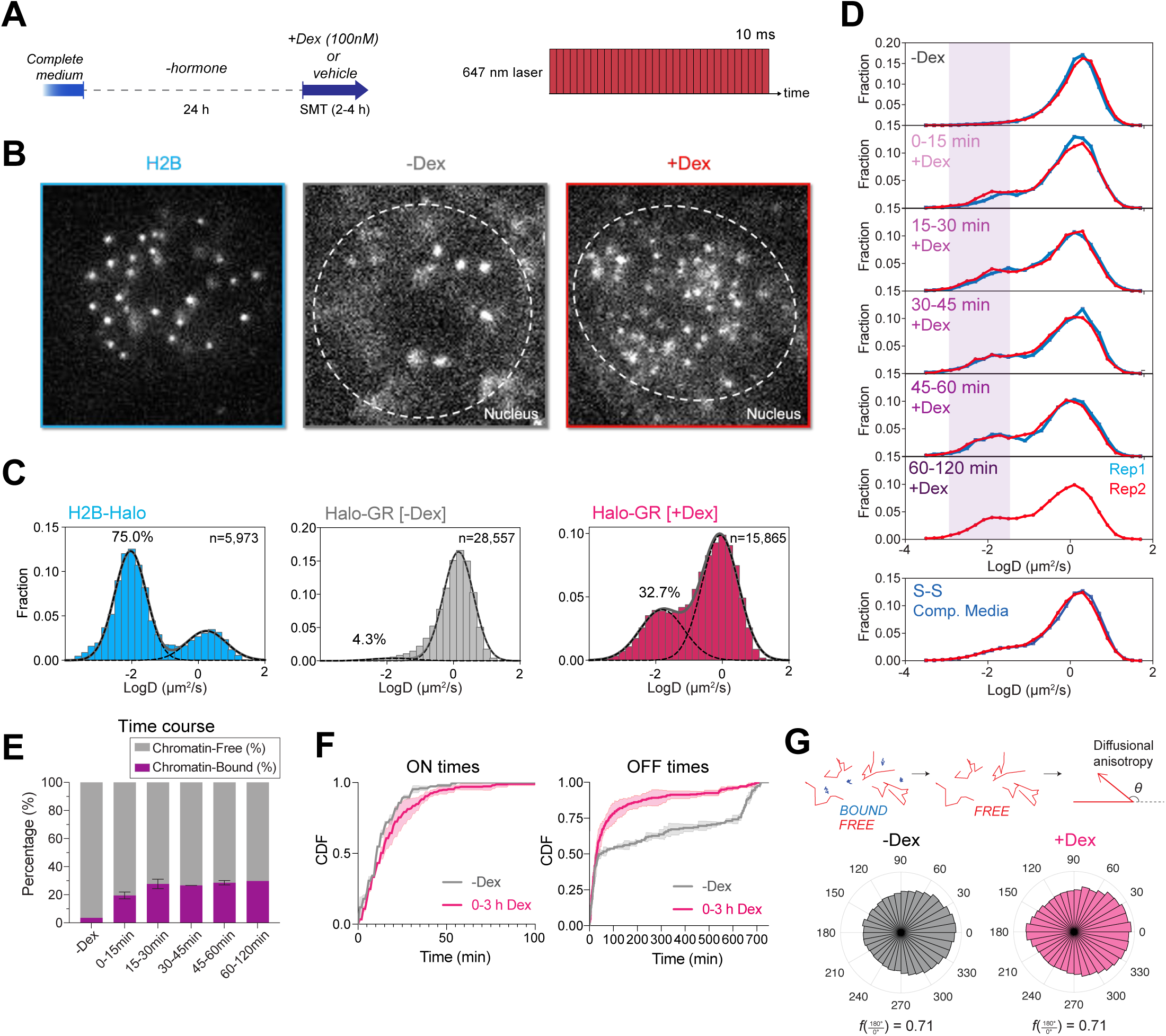
GR can rapidly find its genomic targets in under a minute timescale and activate ERRFI1 transcription. (A) Experimental overview of GR activation by Dex for single molecule tracking (left) and fast tracking (right). (B-C) Representative trajectories in a single nuclei (B) and diffusion coefficient histograms (C) of H2B-Halo and Halo-GR cells in the presence and absence of Dex. (D-E) Diffusion coefficient histograms at different timepoints of Dex treatment (D) with quantification of chromatin-bound and -free percentages (E). (F) CDF plots of *ERRFI1* OFF and ON times in the first 3 hours of Dex treatment compared to -Hormone condition. (G) Schematic illustrating diffusional anisotropy calculation for the freely diffusing population (top), with circular histograms of angular distribution and f(180°/0°) values for -Dex and +Dex conditions (bottom).

To assess whether Dex alters GR’s target search kinetics in the nucleus, we analyzed the diffusional anisotropy of chromatin-free GR trajectories. Diffusional anisotropy quantifies directional bias in consecutive displacements that reflects the geometry of the space sampled by a diffusing molecule (Izeddin et al., 2014). Previously, anisotropic diffusion of nuclear proteins has been associated with spatiotemporal confinement and transient trapping interactions that enhance target search efficiency (Hansen et al., 2020; Ling et al., 2024). However, despite the rapid changes in chromatin-binding frequency upon Dex activation, freely diffusing GR molecules in the nucleus did not display any substantial changes in diffusional anisotropy, indicating that ligand-dependent transcriptional activation is not driven via substantial changes in target search behavior (Fig 3G).

While fast tracking captures global binding frequencies and diffusion-associated behaviors, it does not resolve the lifetimes of chromatin-bound interactions. We therefore complemented these measurements by acquiring 262 ms-frame rate movies (“slow tracking”) to motion-blur freely diffusing molecules and selectively observe chromatin-bound molecules (Fig 4A). Consistent with prior studies, slow tracking revealed ligand-dependent increases in GR chromatin residence times (Fig 4B).

**Figure 4.**
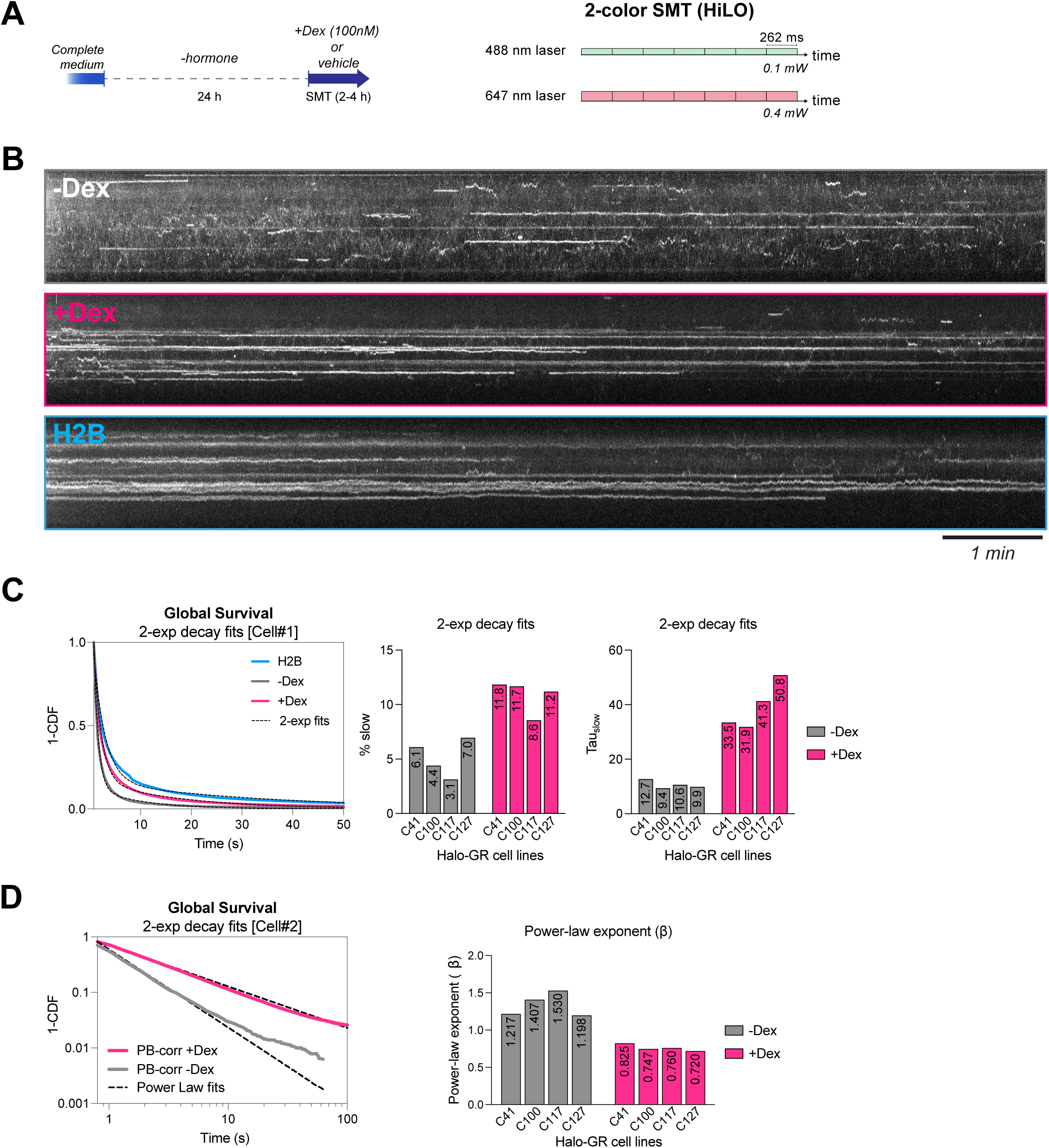
GR shows prolonged chromatin residence times upon Dex induction. (A) Experimental overview of GR activation by Dex for single molecule tracking (left) and slow tracking (right). (B) Representative kymographs for H2B-Halo and Halo-GR (-Dex, +Dex) cell lines. (C) Two-exponential decay fitting of survival curves. Raw curves and fits (left),slow-binding fraction (% slow, middle), and slow residence time (Tau_slow, right) across four stable and endogenously tagged Halo-GR cell lines ± Dex. (D) Power-law fitting of photobleaching-corrected survival curves. Log-log survival plots with photobleaching-corrected ECDF and power-law fits (left), with quantification of the power-law exponent (β) across four Halo-GR cell lines (right).

To quantify global GR residence time distributions, we applied multiple fitting approaches. Double-exponential fits resolved distinct residence time subpopulations and indicated an approximately three-to five-fold increase in the average residence time of the long-lived population across three independent Halo-GR cell lines, from ∼9–13 s in the absence of ligand to ∼32–51 s following Dex treatment, accompanied by an increase in the fraction of long-lived events (from 3-7 to 9-12%) (Fig 4C). In parallel, power-law fits more effectively captured the long-timescale tails of the survival distributions and revealed significant decreases in the power-law exponent β (from 1.2–1.5 to 0.72–0.83) across three independent Halo-GR cell lines (Fig 4D).

Importantly, ligand-dependent increases in GR residence time indicate that dissociation kinetics are dynamically regulated across transcriptional states. We therefore considered whether modulation of GR off-rates may reflect kinetic proofreading processes that enhance transcriptional specificity at target genes.

### Locus-specific tracking reveals dwell-time differences near the GR-target ERRFI1 gene compared to the non-target MYH9 gene

A kinetic proofreading model predicts that differences in transcription factor dwell times contribute to transcriptional specificity, leading to differential transcriptional output at target versus non-target genes. A key feature of these models is that the identity of the activator is sensed at multiple points in the pathway. For example, in a 3-state model both the initial binding state and the dwell time in an intermediate state could be sensitive to activator identity. To directly test this prediction under our experimental system, we performed simultaneous 2-color imaging of GR single molecules and active transcription sites (Fig 5A).

**Figure 5.**
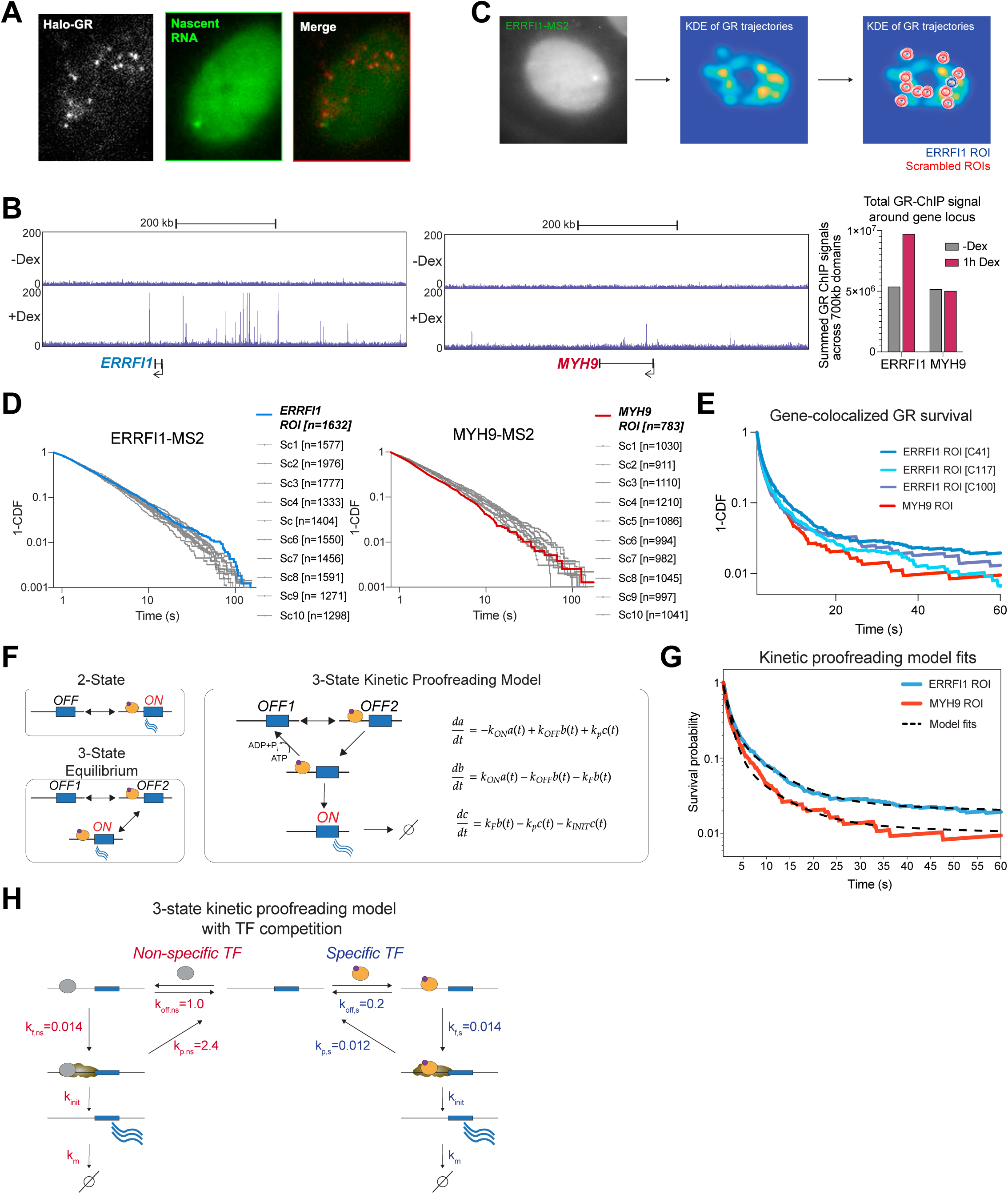
Locus-specific GR tracking shows longer GR survival near its target *ERRFI1* gene compared to non-target *MYH9* gene. (A) Representative snapshot showing two-color imaging of Halo-GR and transcriptionally active *ERRFI1* site. (B) Genome browser tracks of GR-ChIP signals at the 600kb range of *ERRFI1* and *MYH9* genes. (C) Overview of tracking transcriptionally active *ERRFI1* gene to generate *ERRFI1* and scrambled ROIs. (D) 1-CDF curves comparing gene ROI and ten scrambled ROIs for *ERRFI1*-(left) and *MYH9*-MS2 (right) cells. (E) Photobleaching-corrected survival curves for *MYH9* and *ERRFI1* ROIs. (F) Schematic of 2-state, 3-state equilibirum, and kinetic proofreading models with differential equations for the three gene states. (G) GR survival curves at the *ERRFI1* ROI (blue) and *MYH9* ROI (red) fit with a kinetic proofreading mixture model (black dashed lines). (H) A three-state kinetic proofreading model with TF competition. Fitted rate constants (s⁻¹) from (G).

Quantifying transcription factor dwell times at individual regulatory elements is technically challenging due to the limited spatial resolution of optical microscopy. However, several features of our system suggest that locus-proximity–based comparisons may be informative. The *ERRFI1* locus is surrounded by numerous Dex-dependent GR binding sites, whereas *MYH9* lies within a largely GR-depleted region spanning >700 kb by ChIP-seq (Fig 5B), increasing the likelihood that GR molecules detected near these loci reflect enrichment of target-associated or non-target interactions, respectively. Importantly, *MYH9* is also actively transcribed and resides in an open chromatin environment similar to *ERRFI1*, which will not, in theory, prohibit access by GR. Finally, locus identification and analysis by fluorescence microscopy will have the same experimental caveats and limitations for both genes. Thus, we propose that this experimental setup is capable in principle of enriching specific interactions between GR and cognate response elements in the single endogenous gene setting. We therefore asked whether, on average, GR trajectories proximal to the *ERRFI1* locus exhibit different residence times compared with those near the *MYH9* locus.

Because the probability of detecting GR molecules near a given gene locus is low in sparsely labeled Halo-GR population, we acquired a large number of 10-min movies (∼100 movies per cell line), yielding approximately 700–1,000 gene-proximal trajectories per cell line. To minimize filtering-related biases, residence time analyses were performed across multiple radial thresholds and filtering parameters (see Methods). As a negative control, regions of interest (ROIs) defined around diffraction-limited MS2–MCP transcription sites were randomly repositioned within GR-enriched nuclear regions, preserving identical filtering criteria and imaging conditions (Fig. 5C). Compared with scrambled ROI controls, GR survival curves near *ERRFI1* and *MYH9* exhibited subtle but reproducible differences (Fig. 5D). Photobleaching-corrected Halo-GR survival curves from three independent ERRFI1-MS2 cell lines consistently had modest but higher survivals than the MYH9-MS2 cell line (Fig. 5E), demonstrating locus-specific enrichment of GR dwell times at the endogenous target gene.

We next asked whether the observed differences in GR dwell times could be explained by a kinetic proofreading model, and if so, what the underlying rates would imply for transcriptional specificity. While two-state and three-state equilibrium models have been widely used to describe transcription kinetics, both impose an upper limit on specificity set by equilibrium binding affinities (Fig. 5F). In contrast, kinetic proofreading uses irreversible, energy-consuming steps to enhance specificity beyond this limit (Hopfield 1974; Ninio 1975). In this framework, TF bound to a gene can either dissociate (k_off_) or progress to a downstream bound state from which the TF may initiate transcription or be reset to the unbound state by a proofreading step (k_p_) (Fig. 5F). We consider a system where TF binding dynamics reflects both the initial binding and binding as part of the larger complex. Such a model is captured by the following system of ordinary differential equations:

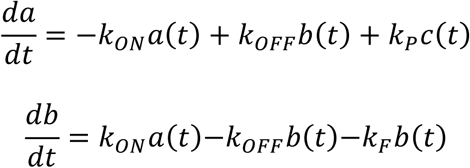

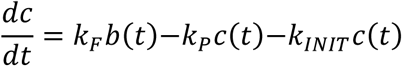

A unique feature of single-molecule tracking is that the observation begins once the molecule is bound. The measurement consists then of observing all possible pathways to return to the unbound state and diffuse away. At steady state during the microscopy experiment, we are thus watching many events that begin in the *b* state and then relax. We can capture this behavior by setting the initial condition to *b[0]=1*. Moreover, *k_ON_* only sets the frequency of events observed in the imaging experiment, which also depends practically on the labeling stoichiometry of the Halo ligand. This *k_ON_* rate therefore does not contribute to the survival curve, computed here:

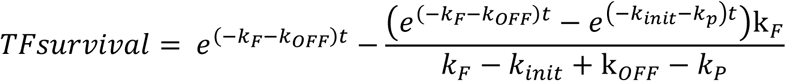

This equation describes the relaxation of the TF and hence the survival curve of a single TF binding a cognate site with no competing sites, an experimental situation which is unrealizable in the nucleus with current microscopy techniques.

In the cellular environment there is competition between GR binding specific and non-specific sites, and the limited spatial resolution of optical microscopy means that each locus-proximal ROI captures a mixture of specific and non-specific binding events. This behavior can be captured by considering competition between specific and non-specific binding according to the following ODE system:

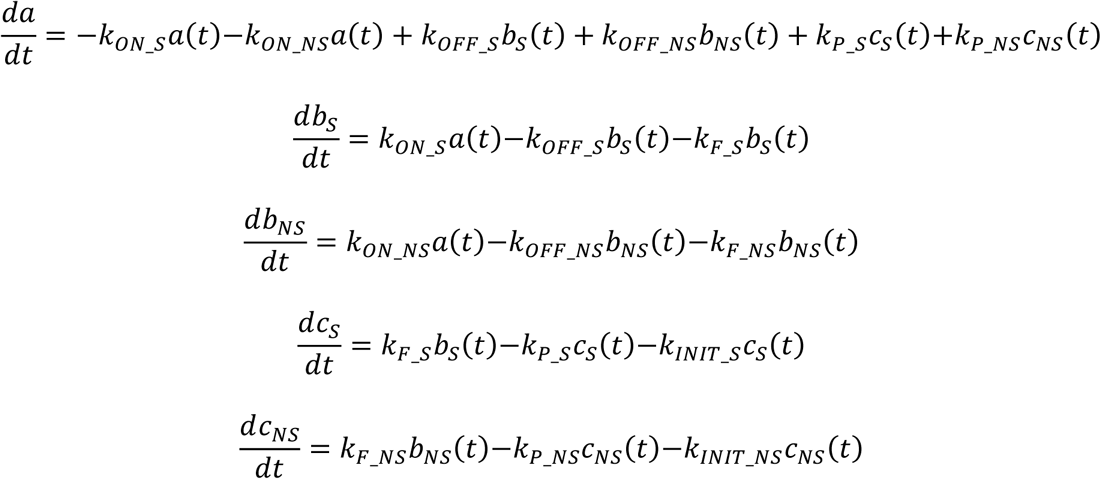

These equations capture the fact that the TF in the unbound state *a* can bind either a specific cognate site or a non-specific site with different kinetics. Now the system is prepared with a mixed initial condition with some fraction *M* bound to the specific site (*b_S_(0) = M*) and some fraction bound non-specifically (*b_NS_(0) = 1-M*). The full solution with these initial conditions is simply:

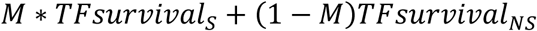

Strikingly, the kinetic proofreading model captured both the *ERRFI1* and *MYH9* survival curves with a single set of rates, differing only in the fraction of specific binding events (Fig. 5G). The fitted rates suggest that the non-specific pathway has ∼5-fold faster dissociation from the initial TF-bound state (k_off,ns = 1.0 s⁻¹ vs. k_off,s = 0.2 s⁻¹), and ∼200-fold faster proofreading rate (k_p,ns = 2.4 s⁻¹ vs. k_p,s = 0.012 s⁻¹) (Fig. 5H). Notably, there is greater discrimination not in the initial binding but in the intermediary state, which we speculate is a complex with other proteins bound to chromatin. The model estimated ∼27% of binding events at the *ERRFI1* ROI are specific compared with ∼14% near *MYH9*, qualitatively consistent with the ChIP-seq enrichment at this locus. Under these fitted rates, even in the presence of a 1,000-fold excess of competing non-specific transcription factors, the model predicts ∼76% transcriptional accuracy at the specific gene (Cho et al., submitted), suggesting that kinetic proofreading provides a plausible mechanism by which the small dwell time differences we observe could be translated into substantial transcriptional specificity.

### A high-throughput imaging-based CRISPR screen identifies regulators of *ERRFI1* and *MYH9* transcriptional bursting under GR activation

To identify regulators of GR–dependent transcriptional activation, we performed a high-throughput, imaging-based CRISPR knockout screen targeting ∼1,100 genes involved in the ubiquitin–proteasome system, cell-cycle, chromatin remodeling, and transcriptional regulation. Nascent transcription of *ERRFI1* and *MYH9* was quantified using a two-color single-molecule RNA FISH assay, in which *ERRFI1* served as a GR-responsive target and *MYH9* as a GR-independent reference gene (Figure 6A). Probe sets targeting the first two introns of *ERRFI1* and the 3′UTR of *MYH9* were used to visualize nuclear transcription sites. Under the high-throughput confocal imaging conditions in 384-well plates, signal-to-noise was insufficient to reliably resolve single dispersed transcripts; consequently, the assay primarily detected discrete nuclear transcription foci corresponding to sites of active transcription.

**Figure 6.**
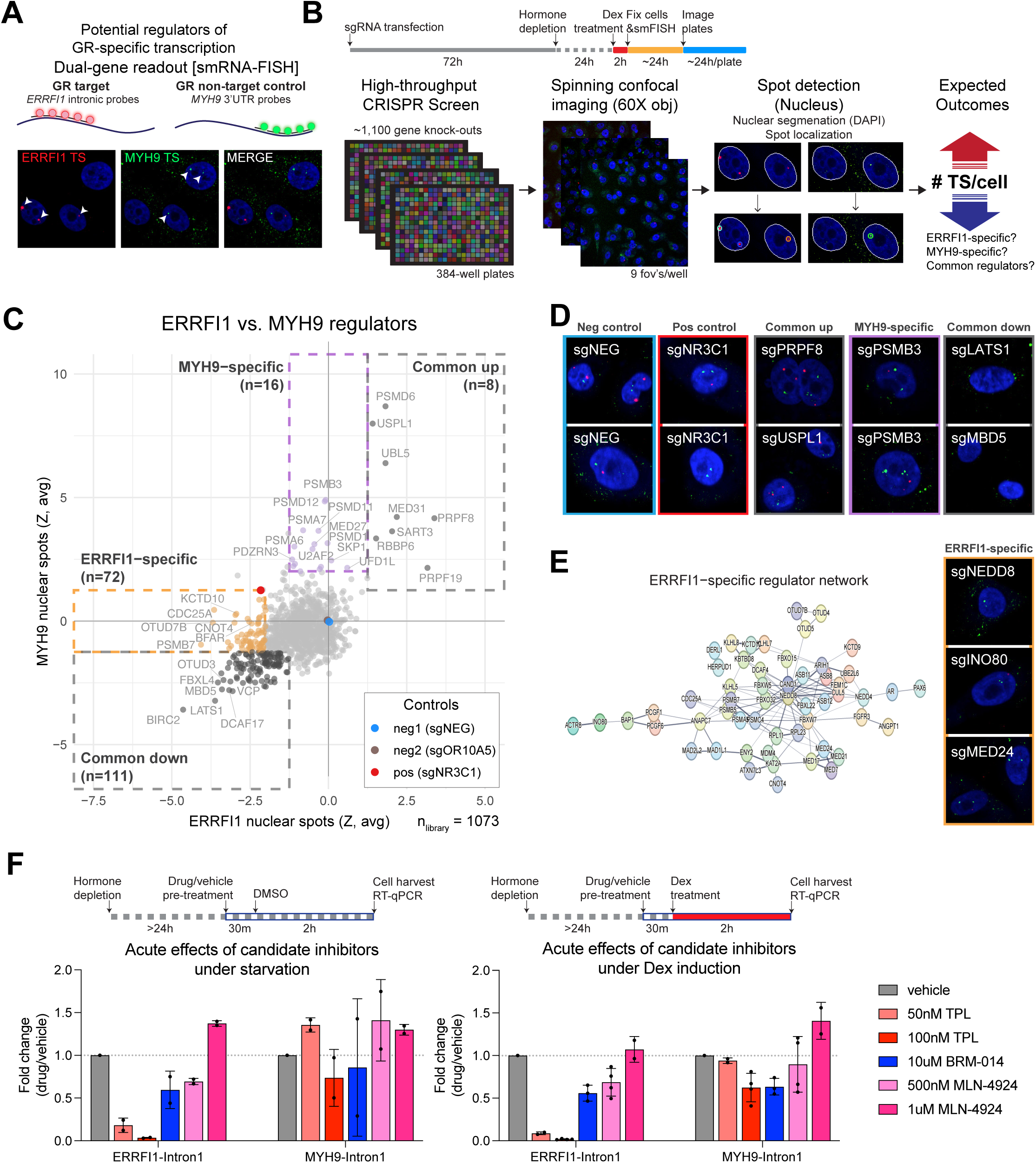
A high-throughput, imaging-based CRISPR screen identifies potential regulators of GR specificity. (A) Screen readout using smRNA-FISH with *ERRFI1*-intronic and *MYH9*-3’UTR probes to detect nascent transcription sites in fixed cells. (B) Screening workflow: sgRNAs targeting ∼1,100 genes are transfected in arrayed 384-well plates, followed by 72-hr knockdown, hormone depletion, dexamethasone treatment, and fixation for smFISH. Plates are imaged by spinning confocal microscopy (60X, 9 fields of view/well), and nuclear spots are quantified per cell via automated DAPI segmentation and spot detection. (C) 2D scatter plot of averaged Z-scores (2 replicates) for *ERRFI1* (x-axis) and *MYH9* (y-axis) nuclear spots, zeroed to the mean of negative controls (sgNEG, sgOR10A5). Hits classified into four categories: “common up” (n=8), “common down” (n=111), “MYH9-specific” (n=16), and “ERRFI1-specific” (n=72). (D) Representative smFISH images for each hit category alongside negative (sgNEG) and positive (sgNR3C1) controls. (E) STRING interaction network of “ERRFI1-specific” hits (left) and representative smFISH images for selected hits (right) (F) RT-qPCR measurement of nascent transcription (intron 1 primers) after 30-min pretreatment with triptolide (TPL; TFIIH inhibitor), BRM-014 (BRM/BRG1 inhibitor), or Pevonedistat (MLN-4924; NEDD8-activating enzyme inhibitor), followed by 2-hr vehicle (left) or 100nM Dex (right) treatment.

The screen was performed in an arrayed format in Cas9-expressing human bronchial epithelial cells, with individual gene knockouts generated by reverse transfection of pooled sgRNAs. Following hormone deprivation, cells were treated with Dex prior to fixation and imaging (Figure 6B). Automated image analysis was used to quantify the number and intensity of nuclear transcription sites for both genes. As biological controls, sgRNAs targeting *NR3C1* (sgNR3C1) served as positive controls for *ERRFI1* but not *MYH9* transcription, whereas scrambled (sgNEG) and *OR10A5*-targeting (sgOR10A5) sgRNAs served as non-cutting and cutting negative controls, respectively (Figure 6C). A lethal sgRNA targeting *PLK1* gene resulted in marked cell loss, indicating high transfection efficiency (>85%). Z-scores were calculated for cell number*, ERRFI1* nuclear spots, *MYH9* nuclear spots, and *MYH9* cytoplasmic spots, with overall concordance observed between two biological replicates (Fig S3). Notably, the distribution of gene knockouts was skewed toward reduced *ERRFI1* nuclear spots (and to a lesser extent *MYH9*), consistent with widespread sensitivity of *ERRFI1* nascent transcription to genetic perturbation and with *ERRFI1* induction being near maximal under Dex treatment.

Joint comparison of *ERRFI1* and *MYH9* nuclear spot z-scores classified the most significant hits into four regulatory categories: (1) ‘common-up’ regulators, whose knockout increased nuclear spots for both *ERRFI1* and *MYH9* (n = 8); (2) ‘common-down’ regulators, whose knockout reduced nuclear spots for both genes (n = 111); (3) ‘MYH9-specific’ regulators, which selectively increased *MYH9* nuclear spots without substantially affecting *ERRFI1* (n = 16); and (4) ‘ERRFI1-specific’ regulators, which selectively reduced *ERRFI1* nuclear spots without substantially affecting *MYH9* (n = 72) (Figure 6C, D).

The common-up regulator group included several proteins involved in RNA processing (PRPF8, PRPF19, SART3, and UBL5), consistent with accumulation of nuclear RNA foci that may reflect retained nascent transcripts due to defects in splicing and/or nuclear export. The common-down regulator group contained genes associated with transcriptional control and general cellular functions; however, many perturbations in this group were accompanied by reduced cell numbers, suggesting that decreased nuclear spots often reflect compromised cell viability or global cellular dysfunction rather than direct transcriptional regulation. Interestingly, multiple proteasome subunit genes (PSMA6, PSMA7, PSMB3, PSMB6, PSMD1, PSMD2, PSMD11, and PSMD12) were present in the MYH9-specific regulator group, indicating a pronounced sensitivity of *MYH9* nuclear spot formation to proteasome perturbation under these conditions.

STRING network analysis of ERRFI1-specific regulators revealed an association network of proteins including NEDD8, a ubiquitin-like protein involved in neddylation; INO80, which encodes the ATPase subunit of an ATP-dependent chromatin remodeling complex; and MED24, a Mediator complex subunit involved in transcriptional regulation during pre-initiation complex assembly (Figure 6E). Disruption of these factors selectively impaired *ERRFI1* nuclear spot formation without comparably affecting *MYH9*, suggesting that ERRFI1-specific regulators may contribute to transcriptional specificity mechanisms that preferentially support robust ERRFI1 induction. Such specificity is consistent with a kinetic proofreading model, in which ATP-dependent regulatory processes selectively amplify productive transcriptional activation at GR-responsive genes.

To functionally test this model on short timescales and minimize indirect effects of chronic gene disruption, we used small-molecule inhibitors targeting ATP-dependent regulatory processes implicated by ERRFI1-specific regulators. Because no selective inhibitors of INO80 or Mediator are currently available, MLN-4924 was used to acutely inhibit neddylation, BRM-014 to inhibit ATP-dependent chromatin remodeling by BRM/BRG1, and triptolide to inhibit the XPB ATPase subunit of TFIIH, which has been previously implicated in transcriptional proofreading (Cho et al., submitted).

Using RT-qPCR to quantify intronic RNA levels as a measure of nascent transcription, we assessed the acute effects of these inhibitors on *ERRFI1* and *MYH9* under hormone starvation and Dex induction (Figure 6F). Triptolide treatment caused a pronounced reduction in *ERRFI1* intron levels under both conditions, whereas *MYH9* intron levels were comparatively resistant, indicating differential sensitivity between the GR-responsive *ERRFI1* gene and the GR-independent *MYH9* gene. In contrast, inhibition of chromatin remodeling or neddylation produced more modest effects on *ERRFI1* induction by Dex, with relatively limited impact on *MYH9* over the same time frame. Together, these results indicate that *ERRFI1* nascent transcription exhibits heightened sensitivity to acute perturbation of ATP-dependent transcriptional processes relative to *MYH9*.

## Discussion

A central question in gene expression is how cells achieve transcriptional specificity to ensure that target genes respond selectively to their specific transcription factors amid the vast excess of non-specific factors in the nucleus. Here, we developed an integrated experimental framework combining single-molecule tracking, live-cell nascent RNA imaging, and high-throughput CRISPR screening to quantitatively link transcription factor binding dynamics with gene-specific transcriptional output at endogenous loci. Using GR as a model inducible transcription factor, we show that transcriptional specificity at its target gene *ERRFI1* arises from modulation of chromatin residence time rather than search kinetics, that this behavior is consistent with kinetic proofreading, and that ATP-dependent processes including neddylation, chromatin remodeling, and TFIIH activity are implicated in this process.

In principle, specificity can be achieved either through ON rates or OFF rates. Through a series of fast and slow tracking under inactive and active GR states, our results suggest that the off-rate modulation, rather than on-rates of transcription factor-chromatin interactions, is the driving kinetic discrimination between target and non-target genes. Moreover, we measure differences in both the off rate from the initial binding state (∼5-fold) and also the intermediate state (∼200 fold). Although classical equilibrium models set an upper bound on regulatory specificity determined by transcription factor concentrations and binding affinities alone (Berg, 2008; Grah et al., 2020; Shelansky et al., 2024; Wong & Gunawardena, 2020; Zoller et al., 2022), data is derived from fitting to a kinetic proofreading model and provides substantial discriminatory power. Interestingly, it is the intermediate binding state which provides greater dynamic range between specific and non-specific binding. Here, gene loci act as dwell-time detectors rather than simple occupancy sensors, discriminating specific from non-specific factors through irreversible, energy-consuming steps (Hopfield, 1974; Ninio, 1975). Indeed, a kinetic proofreading competition model that differed only in the relative fraction of specific binding events at each locus was sufficient to quantitatively capture the survival curves at both *ERRFI1* and *MYH9* ROIs. This finding parallels the observation by Popp *et al*. that transcription factor residence time exerts a stronger influence on burst frequency than concentration (Popp et al., 2021).

A key technical advance of this work is the simultaneous, two-color visualization of endogenous transcription factor single-molecule dynamics and nascent transcription at individual gene loci in living mammalian cells. To our knowledge, this represents one of the first measurements to directly probe the relationship between transcription factor dwell times and transcriptional specificity at endogenous target and non-target genes in their native genomic context. Direct comparison of GR molecules in close proximity to *ERRFI1* and *MYH9* revealed modest but consistent differences in survival distributions, with lower off-rates at the target gene despite both loci residing in transcriptionally active, accessible chromatin. The modest magnitude of these differences between *ERRFI1* and *MYH9*-colocalized GR tracks likely reflects the presence of additional GR target sites within the ROI radius, with GR regulating ∼2,000 genes genome-wide (Figure 1), as well as localization error and the inherent technical limitations of optical microscopy (Tokunaga et al., 2008). Additionally, our 2D tracking and 10-minute acquisition windows impose temporal limits on observed GR binding events, and because the MS2 system marks only sites of active transcription, locus-resolved measurements are inherently biased toward the transcriptionally active state.

Our high-throughput, imaging-based CRISPR screen and subsequent pharmacological validation experiments illuminate the molecular machinery that may underlie kinetic proofreading at GR-responsive loci (Figure 6). Among the *ERRFI1*-specific regulators identified (i.e. gene knock-out that selectively disrupted *ERRFI1* but not *MYH9* transcription), two classes of ATP-dependent processes emerged as particularly noteworthy. First, proteins involved in the neddylation pathway, including NEDD8 and CAND1, were required for full *ERRFI1* induction. Neddylation primarily activates cullin-RING E3 ubiquitin ligases, broadly regulating ubiquitin-proteasome-dependent turnover of hundreds of substrates (Petroski & Deshaies, 2005), though interestingly, direct neddylation of non-cullin transcription factors such as p53 and E2F-1 has also been reported (Loftus et al., 2012; Xirodimas et al., 2004). Consistent with neddylation pathway’s broad role in ubiquitin-proteasome system, the proteasome has been shown to be involved in increasing off-rates of nuclear receptors, including GR and ER, at their target sites (Reid et al., 2003; Stavreva et al., 2004; Stenoien et al., 2000). The second group was ATP-dependent chromatin remodelers, such as INO80. ATP-dependent chromatin remodelers consume energy to reposition nucleosomes or evict histone octamers, where intriguingly, remodelers can also displace transcription factors on the templates undergoing nucleosome remodeling in an ATP-dependent manner (Li et al., 2015; Nagaich et al., 2004). Furthermore, an independent study from our laboratory found TFIIH as a promising candidate for regulating kinetic proofreading rate (Cho et al., submitted). Like other screen hits, acute inhibition of the XPB ATPase subunit of TFIIH by triptolide selectively impaired *ERRFI1* nascent transcription while largely sparing *MYH9*. Together, these ATP-dependent processes may constitute kinetic proofreading processes that discriminate between specific and non-specific GR transcription factor interactions, returning the promoter to a vacant state when the incorrect factor is bound.

In summary, our study integrates single-molecule GR kinetics with nascent transcription imaging at endogenous target and non-target gene loci to address transcriptional specificity. Collectively, our findings support a model in which transcriptional specificity is an emergent property of ATP-dependent molecular processes that shape stochastic gene transcription.

## Materials and Methods

### Cell culture and treatments

Human bronchial epithelial cells (cell lines derived from HBEC3-kt; ATCC, CRL-4051) were maintained at 37°C in a humidified incubator with 5% CO_2_. Cell lines used in this study can be found in Supplementary Table 1. Cells were cultured in Airway Epithelial Cell Basal Medium (ATCC, PCS-300-030) supplemented with 1% penicillin/streptomycin and Bronchial Epithelial Cell Growth Kit (ATCC, PCS-300-040) containing HLL Supplement, L-glutamine, Extract P, and Airway Epithelial Cell Supplement (“complete medium”). For hormone depletion (“-Hormone” or “starved” conditions), cells were washed one to three times and incubated for >24 h in ATCC basal medium supplemented with only HLL Supplement, L-glutamine, and 1% penicillin/streptomycin.

For the RNA-seq experiment (Fig. 1), hormone-depleted cells were treated with 100 nM dexamethasone (Dex; Sigma-Aldrich, D4902-25MG), 100 nM LG268 (Tocris, Cat. No. 5920), 100 nM AM580 (Tocris, Cat. No. 0760), or 1 µM rosiglitazone (RGZ; Cayman Chemical, Cat. No. 71740) for 2 h or 4 h, as indicated. For inhibitor perturbations (Fig. 6,7), hormone-depleted cells were pre-treated for 30 min with triptolide (TPL; 50 nM or 100 nM) (Sigma-Aldrich, T3652), BRM-014 (10 µM) (MedChemExpress, HY-119374), or MLN-4924 (500 nM or 1 µM) (Cell Signaling, #85923), followed by 2 h treatment with Dex (100 nM) or vehicle in the continued presence of inhibitor, as indicated. All compounds were prepared as stock solutions in DMSO, except Dex, which was dissolved in ethanol, and were diluted into culture medium immediately prior to treatment (final ethanol/DMSO concentration ≤0.1% v/v).

### Stable Halo-GR cell line generation using lentiviral transduction

Stable Halo-GR cell lines were generated using lentiviral transduction. The coding sequence for glucocorticoid receptor alpha (GRα) was cloned into the N-terminal HaloTag vector pFN21A to produce a HaloTag-GRα fusion construct. The Halo-GRα cassette was then subcloned into a lentiviral transfer plasmid, and lentiviral particles were used to transduce cells for stable genomic integration. Lentivirus was produced in HEK293T cells cultured in high-glucose DMEM (Gibco, Cat. No. 10569010) supplemented with 10% fetal bovine serum (FBS) and 1% penicillin/streptomycin. HEK293T cells were seeded on a 6-cm dish at approximately 90% confluency and transfected with 15 µg of lentiviral transfer vector along with 0.75 µg each of Tat, Rev, and Gag/Pol, and 1.5 µg of VsvG packaging plasmids using Lipofectamine 3000 (Invitrogen, Cat. No. L3000015). Viral supernatants were collected over 3 days and concentrated using PEG-it Virus Precipitation Solution (SBI, LV825A-1) and filtered through a 0.45 µm filter. Low-passage HBECs were transduced with lentivirus in the presence of 10 µg/mL polybrene (Sigma-Aldrich, TR-1003-G). Transduced cells were expanded and single-cell cloned by fluorescence-activated cell sorting (FACS) on an MA900 cell sorter (Sony Biotechnology) using “Single 60” sort mode into 96-well plates.

### Endogenous HaloTag knock-in at NR3C1 (Halo-GR) using CRISPR–Cas9 genome engineering

To generate an endogenous Halo-tagged glucocorticoid receptor (Halo-GR), a HaloTag coding sequence was inserted at the NR3C1 translational start site (ATG) by homology-directed repair, resulting in an N-terminal HaloTag fusion at the endogenous NR3C1 locus (Fig. 2A). The sgRNA sequences, donor design, and linker sequences are provided in Supplementary Table 2. CRISPR–Cas9 genome engineering was performed in low-passage (<3-5 passages) HBECs using electroporation with the Neon Transfection System (Invitrogen, MPK5000; 1230V, 10 ms pulse width, 4 pulses). Cells were electroporated with two plasmids: (i) a plasmid expressing Cas9 fused to mCherry along with a single guide RNA (sgRNA), and (ii) a donor template plasmid containing 500-bp homology arms flanking the desired insertion. At 72 h post-electroporation, cells were enriched by FACS for mCherry expression and HaloTag ligand labeling. Upon expansion of mCherry-positive cells, single cells were labelled with JF646 HaloTag ligand and sorted based on JF646 expression on an MA900 cell sorter (Sony Biotechnology) in 96-well plates. Clonal lines were expanded and validated by genomic PCR and western blotting.

For acute depletion of GR in homozygously tagged Halo-GR cell lines, cells were treated with HaloPROTAC3 (Promega, GA3110) at 500 nM for the indicated durations. For experiments combining GR depletion with Dex induction, cells were hormone-depleted for 18 h, treated with HaloPROTAC3 for 6 h, and then stimulated with Dex for the final 2 h in the continued presence of HaloPROTAC3 (Fig. 2C–D). GR depletion was confirmed by western blotting.

### Single molecule tracking sample preparation, data acquisition, and analysis

#### Cell preparation and dye labeling for SMT

For Halo-GR single-molecule tracking (SMT), cells were seeded in 4- or 8-well Lab-Tek chambered coverglass (Thermo Scientific, 155382 or 155409) and hormone-depleted for >24 h. Cells were then treated with 100 nM Dex or vehicle control prior to imaging. HaloTag fusion proteins were labeled with 2.5–5 pM JFX650 for 15 min at 37°C, followed by >5 washes with culture medium to remove unbound dye. H2B-Halo cells were labeled identically except using 0.5–0.75 pM JFX650 (Donovan et al., 2024).

#### Image acquisition for single-molecule tracking

Imaging was performed on a custom-built HiLO microscope (Mehta et al., 2018). This custom-built microscope from the CCR, LRBGE Optical Microscopy Core facility is controlled by μManager software (Open Imaging, Inc., San Francisco, CA.), equipped with an Okolab stage top incubator for temperature control (37°C) with an objective heater, a 100X 1.49 numerical aperture objective (Olympus Scientific Solutions, Waltham, MA), 647nm and 488nm lasers (Coherent OBIS, Santa Clara, CA), and EM-CCD camera (Evolve 512 Delta, Photometrics).

For slow tracking, GFP and JFX650 channels were acquired simultaneously with an exposure time of 262 ms per frame for both lasers, and a time-lapse interval of 262 ms for a total movie duration of 10 min, with laser powers of 0.3-0.4 mW (647 nm) and 0.01-0.03 mW (488 nm). For cell lines with low GFP-MS2 expression (e.g., cJK100), a pulsed 488-nm laser scheme (1 s on, 2 s off) was employed to minimize photobleaching.

For fast tracking, JFX650 images were acquired simultaneously with an exposure time of 10 ms and a time-lapse interval of 10 ms for a total movie duration of 30 sec, with a laser power of 15-16 mW (647 nm).

#### Single-molecule tracking analysis

We used the custom-made software TrackRecord (Mazza et al., 2013) in MATLAB (The MathWorks, Inc., Natick, MA). To analyze each time series, data were filtered using top-hat, Wiener, and Gaussian filters. A region of interest (ROI) was defined to encompass the nuclei based on the nuclear GFP staining, then nuclear particles were detected, fitted to two-dimensional Gaussian function for subpixel localization, and finally tracked using a nearest neighbor algorithm (Presman et al., 2017).

For slow tracking, the Halo-GR tracking parameters were as follows: window size for particle detection 3 pixels, maximum frame to frame displacement of 3 pixels, shortest track 3 frames, and gaps to close. MS2-labeled nascent RNA spots were tracked using the same parameters as Halo-GR molecules, but with more permissive gap-closing parameters, since MS2 spots exhibit minimal displacement over the 10-min imaging window.

For fast tracking, the parameters were as follows: window size for particle detection 3 pixels, maximum frame to frame displacement of 8 pixels, shortest track 5 frames, and gaps to close 0.

#### Fast tracking (Diffusion coefficient) analysis

Halo-GR trajectories from fast-tracking data were analyzed using the MSD analyzer in MATLAB (Tarantino et al., 2014). For each trajectory, the time-averaged mean square displacement (TA-MSD) was computed, and the diffusion coefficient (*D*) was estimated by linear regression of the first 5 time-lags of the TA-MSD curve:

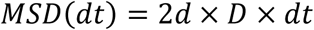

where *d* is the number of dimensions (*d* = 2). Trajectories with R² <0.8 were excluded.

#### Slow tracking (1-CDF) analysis

Halo-GR trajectories from slow-tracking data that were shorter than 3 frames (0.786 s) were excluded, and survival curves were constructed as the complement of the cumulative distribution function (1-CDF), binned at 262 ms. 1-CDF curves were fit to a double exponential decay model:

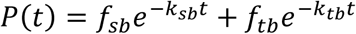

where k_sb and k_tb correspond to dissociation rates for stable- and transient-binding events, respectively, and 1 = f_sb + f_tb for the two components. The apparent k_sb and k_tb values are affected by photobleaching and chromatin movement. To correct for this bias, we used apparent dissociation rates of H2B imaged under the same conditions as described previously (Hansen et al., 2017). The corrected residence times for stable (τ_sb) and transient (τ_tb) binding were calculated as follows:

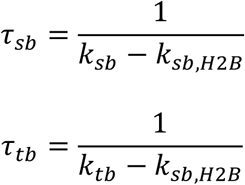

For power-law fitting of 1-CDF curves, we followed the ECDF-based approach described previously (Garcia et al., 2021). The empirical survival function was computed from unbinned dwell times, and corrected for photobleaching by dividing by:

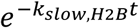

The photobleaching-corrected survival was fit to a power-law model:

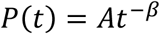

where A is a normalization constant and β is the power-law exponent.

#### Two-Color Imaging and ROI-Based Colocalization Analysis

Because sufficient statistical power at individual gene loci required the acquisition of many imaging sessions across multiple days and cell lines, the following standardization procedures were implemented to ensure consistency of the two-color imaging data. On each imaging day, to ensure the alignment of the images collected in green and far red, which are obtained on separate cameras, we first collected images of 0.2 mm TetraSpeck beads (Fisher Scientific, T7280) in both channels. Five to eight matching beads were then selected, and their positions were used with MATLAB’s built-in function fitgeotrans to find an affine transformation between the two channels. The resulting transformation was then applied to the green channel prior to selecting the ROI. Application of the transformation to the bead images themselves indicated that the error in alignment was ∼1.4 pixels (0.2 µm). Next, H2B-Halo cells labeled under identical conditions were imaged as a reference standard each day or every other day (Donovan et al., 2024). Because H2B is stably bound to chromatin, H2B-Halo served as an internal control for both photobleaching kinetics and day-to-day variation in laser power and alignment. Residence-time survival curves were corrected using the H2B-Halo photobleaching decay (see “Photobleaching Correction” below).

To measure GR residence time at specific gene loci, an ROI-based colocalization analysis was developed using the standardized two-color SMT data (Fig. 5C). For each cell, the MS2-labeled transcription site (marking the *ERRFI1* or *MYH9* locus) was tracked in the 488-nm channel, and a circular ROI was defined around the MS2 spot position at each frame. In the 647-nm channel, a kernel density estimate (KDE) of Halo-GR trajectory positions was computed using MATLAB’s *mvksdensity* function (bandwidth = 5 pixels) on a 256 × 256 pixel grid. To avoid biasing the density map toward long-lived molecules, only one localization per trajectory (the mean *x* and *y* position) was used for KDE computation.

A Halo-GR trajectory was classified as gene-colocalized if, at every frame, the molecule fell within the ROI radius of the MS2 spot position (strict colocalization criterion). Residence times of gene-colocalized GR molecules were then extracted and analyzed as described above. To control for locus-specific effects, ten scrambled ROIs were generated per cell by sampling random positions from the KDE density map using weighted random sampling; positions with higher GR density were thus selected with proportionally higher probability. Each scrambled ROI preserved the frame-by-frame displacement of the MS2 spot but was translated to the newly sampled position. Scrambled ROIs were constrained to be non-overlapping with one another and with the MS2 ROI. For each scrambled ROI, the same strict colocalization criterion was applied, and residence times were extracted. The gene-locus ROI survival curve was then compared with the ten scrambled ROI survival curves to assess whether GR exhibits prolonged binding specifically at the target gene (Fig. 5D).

To generate photobleaching-corrected survival curves for gene-colocalized GR molecules (Fig. 5E), dwell times from all gene-locus ROI colocalization events were pooled across imaging sessions and compiled into a single 1-CDF survival curve. To correct for photobleaching and chromatin motion, H2B-Halo 1-CDF curves acquired under identical imaging conditions were first computed separately for each imaging date and then combined as a weighted average, with weights proportional to the number of GR trajectories acquired on the corresponding date. The weighted-average H2B 1-CDF was fit to a triple-exponential decay model, and the slowest decay rate (𝑘_𝑠𝑙𝑜𝑤,𝐻2𝐵_) was extracted as the photobleaching correction factor as described previously (Garcia et al., 2021). The corrected survival function was then calculated by dividing the gene-colocalized GR 1-CDF by ⅇ^−𝑘𝑠𝑙𝑜𝑤,𝐻2𝐵^^𝑡^.

### RNA extraction and RT–qPCR

Total RNA was purified using the RNeasy Mini Kit (Qiagen, 74104) and treated with TURBO DNA-free Kit (Thermo Fisher, AM1907) to remove genomic DNA. cDNA was generated from DNase-treated RNA using the ProtoScript II First Strand cDNA Synthesis Kit (NEB, E6560L) with the manufacturer-supplied random primer mix. RT–qPCR was performed using iQ SYBR Green Supermix (Bio-Rad, 1708880) on a CFX96 Touch Real-Time PCR Detection System (Bio-Rad). Gene expression was quantified using the ΔΔCt method with ACTB as a housekeeping control. Primer sequences are listed in Supplementary Table 3.

### RNA- and ChIP-seq Analysis

Early-passage wild-type HBECs were cultured in complete medium and then hormone-depleted for 24 h prior to treatment with Dex, LG268, AM580, or RGZ for 2 h or 4 h, as indicated. RNA-seq was performed with three biological replicates per condition. Total RNA was extracted and DNase-treated as above. rRNA-depleted libraries were prepared using the NEBNext rRNA Depletion Kit v2 (Human/Mouse; NEB, E7405) followed by the NEBNext Ultra II Directional RNA Library Prep Kit for Illumina (NEB, E7760) and NEBNext Multiplex Oligos for Illumina (96 Unique Dual Index Primer Pairs; NEB, E6440). Libraries were quality-controlled using TapeStation and sequenced on an Illumina NextSeq 2000 using a P3 200-cycle kit. For RNA-seq analysis, adapter trimming was performed with Trim Galore (v 0.6.7) with FastQC enabled. Reads were aligned to the human genome (GRCh38) using HISAT2 (v 2.2.1) with reverse-stranded settings (RF). Gene-level read counts were obtained using HTSeq-count (v 2.0.4) with strandedness set to “reverse” and an GENCODE v38 comprehensive gene annotation. Differential expression was calculated with DESeq2 using significance threshold of adjusted p-value < 0.05. ChIP sequencing was performed and analyzed as described previously (Johnson et al., 2021).

### Western blotting

Whole-cell lysates were separated by SDS–PAGE 4-12% Bis-Tris gel (Invitrogen, NP0322BOX) and transferred to 0.2 µm nitrocellulose membranes (Bio-Rad, #1704270) using Trans-Blot Turbo Transfer system (Bio-Rad). Membranes were probed with anti-GR (Santa Cruz, sc-393232 X; 1:10,000 dilution), anti-Halo (Promega, G9211; 1:1,000 dilution), anti-β-Actin (Sigma-Aldrich, A2228; 1:10,000 dilution), and anti-H2B (Abcam, ab52484; 1:,2,000 dilution) antibodies, followed by HRP-conjugated secondary antibodies (1:5,000 dilution) and chemiluminescent detection (Thermo Scientific, 34095).

### Single-molecule fluorescence *in situ* hybridization (smFISH)

smFISH was performed as described previously, using the same probe sets targeting *ERRFI1* introns (Quasar 670) and the 3′UTR of *MYH9* (Quasar 570) (Patange et al., 2022; Wan et al., 2021). Probe sequences are provided Supplementary Table 4. Briefly, cells growing on glass coverslips (18 mm, #1.5 thickness; Electron Microscopy Sciences, 72222-01) in 12-well plates were fixed for 10 min in 4% PFA in PBS, washed twice for 10 min each in PBS, and permeabilized in 70% ethanol at 4°C for at 1-3 days. Hybridization was carried out in hybridization buffer (10% dextran sulfate, 10% formamide, 2× SSC) and freshly prepared wash buffer (10% formamide, 2× SSC). Permeabilized cells were equilibrated in wash buffer for 5 min at room temperature, then incubated face-down on 50-µL drops of probe diluted to 100 nM in hybridization buffer in a humidified chamber at 37°C for 4–16 h in the dark. Coverslips were subsequently washed twice for 30 min each at 37°C with wash buffer, rinsed briefly in 2× SSC, washed for 5 min in PBS at room temperature, and mounted on glass slides using ProLong Gold Antifade Mountant with DAPI (Invitrogen).

Samples were imaged on a custom Rapid Automated Modular Microscope (RAMM; ASI Imaging) equipped with a 40×/1.4-NA oil-immersion objective (Zeiss), an ORCA-Flash4 V2 sCMOS camera (Hamamatsu), a SpectraX light engine (Lumencor), and a quad-bandpass filter set (DAPI/GFP/Cy3/Cy5; Chroma Technology). Additional single-band emission filters for Cy3 (605/70) and Cy5 (700/75; Semrock) were housed in a high-speed filter wheel (ASI Imaging). Automated acquisition was controlled through MicroManager (Edelstein et al., 2010). Excitation wavelengths and camera exposures were as follows: 395 nm/50 ms (DAPI), 550 nm/400 ms (Cy3/Quasar 570), and 640 nm/400 ms (Cy5/Quasar 670). Z-stacks were acquired at 0.5-µm step size to capture the full cell volume (∼21 slices for HBECs) and converted to maximum-intensity projections. Image analysis was performed using a custom Python pipeline (https://github.com/Lenstralab/smFISH_analysis.git).

### Arrayed CRISPR-KO Ubiquitin Screen

A library of chemically modified synthetic sgRNA oligos targeting 1159 genes involved in the ubiquitin and proteasome pathways (Table S5 for list of genes and sgRNA sequences) was purchased from Synthego. Each gene was targeted by 1-3 oligos bioinformatically designed by Synthego to produce staggered cuts in a 50 – 100 bp exonic span close to the transcriptional start site of the gene. The 3 oligos per gene were delivered as a dried pool in each well of a 96-well library. We resuspended the sgRNA oligos in ddH2O, and then transferred and reformatted them in 384-well ECHO-compatible plates. A Beckman ECHO525 was used to spot 0.08 pmoles/well of the sgRNA library in a 325 nl volume at the bottom of empty Revvity imaging 384-well plates. The library was spotted in columns 2 – 21. We spotted the same volumes and amount of control sgRNA oligos in columns 23 and 24: sgNEG (Negative Control, Scrambled sgRNA#2), sgOR10A5 (negative non-expressed gene control), sgPLK1 (Positive transfection control), sgNR3C1 (encoding for GR, biological positive control). The plates were air dried under a laminar flow cell culture hood, sealed, and then stored at -20C until the day of the transfection.

On the day of transfection, plates were thawed and equilibrated at RT for at least 30 min. Plates were then spinned for 5 min at 1400 rpm in a tabletop centrifuge and unsealed. 20 μl of “complete medium” described above without penicillin/streptomycin containing 50 nl of RNAiMax Lipofectamine transfection reagent complex (ThermoFisher, 13778-150, lot 2539523) were added to each well using a ThermoFisher MultiDrop Combi liquid dispenser, and incubated for 30 min at RT. 20 μl per well of a suspension of 1,100 HBEC-Cas9 cells in “complete medium” without penicillin/streptomycin was added to the sgRNA oligos and transfection reagents complexes. The final volume was 40 μl and the final sgRNA oligo concentration was 2 nM. Plates were incubated for 30 min at RT, and then transferred to 37C in a humidified and CO2-controlled incubator. At 72 h post-transfection, media was removed using ‘Gentle Spin’ on BlueWasher (BlueCatBio), and 40 µL “-Hormone” media was added using the LOW speed setting on MultiDrop liquid dispenser for hormone starvation for 24 h. 10 µL 500nM Dex were added to all wells for 2h before cells were fixed. The screen was run in 2 biological replicates.

#### High-Throughput HCR RNA FISH Staining

384-well imaging plates were fixed by adding 40 µL 8% paraformaldehyde (PFA) (4% final) for 15 min. Fixed cells were washed 2x in DPBS for 7 min at RT, and permeabilized in 80uL 70% ethanol for >1 h at -20°C. Permeabilized cells were washed 1X DPBS for 10 min and incubated wash buffer (10% Formamide, 2X SSC) for 30min at RT. Primary probes (50 nM each; ERRFI1 intron probe set labeled with Q670 and MYH9 3′UTR probe set labeled with Q570) were hybridized for 20–24 h at 37°C in 10uL hybridization buffer (10% dextran sulfate, 10% formamide, 2× SSC) in a humidified chamber. Plates were washed twice for 30 min in wash buffer (10% formamide, 2× SSC), followed by 10 min washes in 2× SSC and 1× DPBS. Nuclei were stained with DAPI (2.5 ng/µL) for 30 min and washed once in 1× DPBS. Plates were stored at 4C until ready for imaging.

#### High-content imaging and spot detection

Images were acquired on a Yokogawa CV8000 high-throughput spinning disk confocal microscope. Imaging plates were excited with 405, 488, 561, or 640 nm laser in separate exposures using a near IR autofocus mechanism, a 405/488/561/640 nm excitation dichroic mirror, a 63X plan apochromat water objective (NA 1.2), a 561 nm emission dichroich splitting mirror, and 2 sCMOS cameras (2048 X 2048 pixels, bin 2X2, pixel size 216 nm). For each field of view (FOV), we acquired a 8 μm z-stack at 0.5 μm intervals in each channel. Images were corrected on the-fly using Yokogawa proprietary software to compensate for background, illumination vignetting, geometric channel registration, and chromatic aberrations. Corrected z-stacks were then maximally projected on the fly and saved as TIFF files (mrf/tiff format). We acquired 9 FOVs per well.

#### High-Throughput Image Analysis

Images acquired on the CV8000 were imported and analyzed in Revvity Signals Imaging Artist (SiMA, version 1.4.3). Briefly, nuclei were segmented using the DAPI channel and then used as seeding object for cell body segmentation using the cytoplasmic background signal in the DAPI channel. Cells whose cell body was touching the image border were excluded from the analysis. A spot finding algorithm was used to find HCR RNA FISH signals in the 561 and 640 channels in the nucleus and cytoplasm regions, respectively, and spot fluorescence intensity was measured. Single cell results for each calculated cellular feature were mean aggregated over the cell population in a well. Well-level results were exported as tabular format as separated text files.

#### Screen Statistical analysis

Well-level results from the SiMA image analysis of the sgRNA KO screen against the Ubiquitin/Proteasome library were analyzed using using R 4.3.3 and the cellHTS2 package 2.66.0 (Boutros et al., 2006). Briefly, per well raw measurements for all the features calculated by SiMA were normalized on a per plate basis using the median value of the sgRNA library treatments and the Z-score method. All the per plate normalized values in different library plates originated from a single biological replicate were further standardized using a robust version of the Z-score. The Z-score values for the same well and plate combination in different biological replicates were then averaged to obtain a Mean Z-score per gene KO and per calculated feature. To account for residual offset of the negative control distributions from zero, the Mean Z-scores were re-centered by subtracting the mean Z-score of the two non-targeting negative controls (sgNEG and sgOR10A5) for each readout independently. Gene knockouts associated with significant cell death (mean cell count < 30 across replicates) were excluded from downstream hit calling, yielding a final set of 1,073 gene knockouts for analysis.

Hits were classified into four categories using the control-centered Z-scores for ERRFI1 and MYH9 nuclear spot counts. ERRFI1-specific regulators were defined as gene knockouts with a centered ERRFI1 Z-score < −2 and a centered MYH9 Z-score within ±1.25 of zero. MYH9-specific regulators were defined as knockouts with a centered MYH9 Z-score > 2 and a centered ERRFI1 Z-score within ±1.25 of zero. Common regulators were identified as knockouts for which both centered Z-scores exceeded 1.25 in the same direction: common up-regulators (both Z-scores < −1.25) and common down-regulators (both Z-scores > 1.25). Gene knockouts falling outside these thresholds were classified as non-hits.

### Live-cell, nascent RNA imaging and analysis

Cells were seeded in 96-well glass-bottom plates (Cellvis, P96-1.5H-N) and imaged on a Yokogawa CV7000 high-throughput spinning disk confocal microscope maintained at 37°C with 5% CO_2_. The optical configuration consisted of a 405/488/561/640-nm excitation dichroic mirror, a 63× plan-apochromat water-immersion objective (NA 1.2), a 561-nm emission dichroic beam splitter, and two sCMOS cameras (2560X2160 pixels; 2 × 2 binning; effective pixel size of 216 nm). Cells were excited with a 488-nm laser at 25% power. Time-lapse movies were acquired over 12 h at 100-s intervals. For each time point, eight fields of view (FOVs) were captured, with each FOV comprising a 5-µm z-stack acquired at 0.5-µm step size. Drug treatments (Dex and/or HaloPROTAC3) were applied as indicated. Images were corrected in real time using Yokogawa proprietary software to compensate for background noise, illumination unevenness, inter-channel geometric registration, and chromatic aberrations. Corrected z-stacks were maximum-intensity projected on the fly and saved as TIFF files.

Nuclear segmentation and rotational registration of individual nucleus ROIs were performed using the HiTIPS analysis pipeline (Keikhosravi et al., 2024). Registered single-nucleus images were then processed with a custom IDL script (*localizeApp.sav*) to detect and track transcription sites across the time-lapse series. Transcriptional ON and OFF states were subsequently assigned by fitting hidden Markov models (HMMs) to the fluorescence intensity time traces of individual transcription sites using a second custom IDL script (*fcsApp.sav*).

## Supporting information

Supplemental Figures_combined

Supplemental Tables_combined

