## Supplemental Figures_combined for "Kinetic proofreading as a mechanism for transcriptional specificity in living human cells"

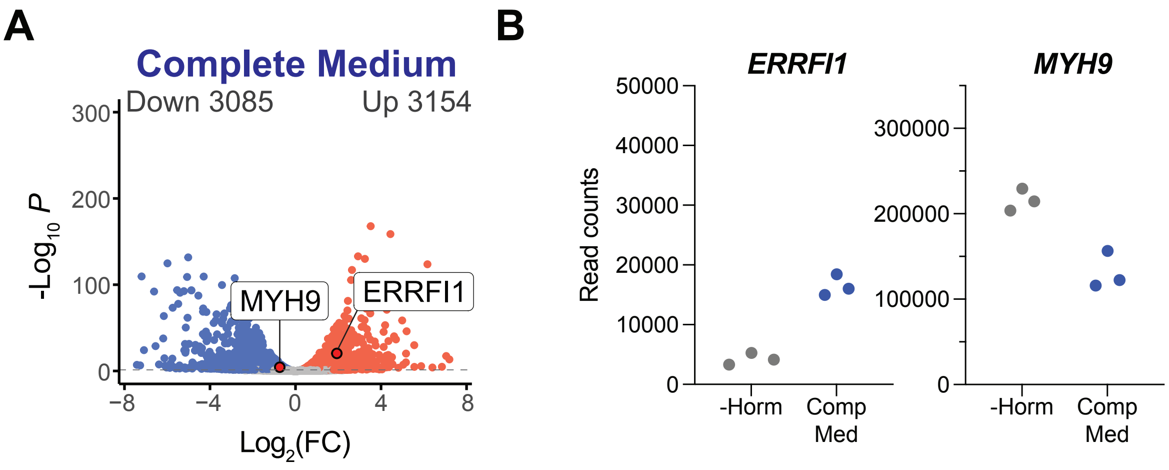


**Figure S1. *ERRFI1* and *MYH9* expression in complete medium.**
(A) Volcano plot showing differentially expressed genes in complete medium versus hormone-deprived conditions (dashed lines indicate adj p-val threshold of 0.05).
(B) Read counts for *ERRFI1* and *MYH9* in hormone-deprived (-Horm) and complete medium conditions.

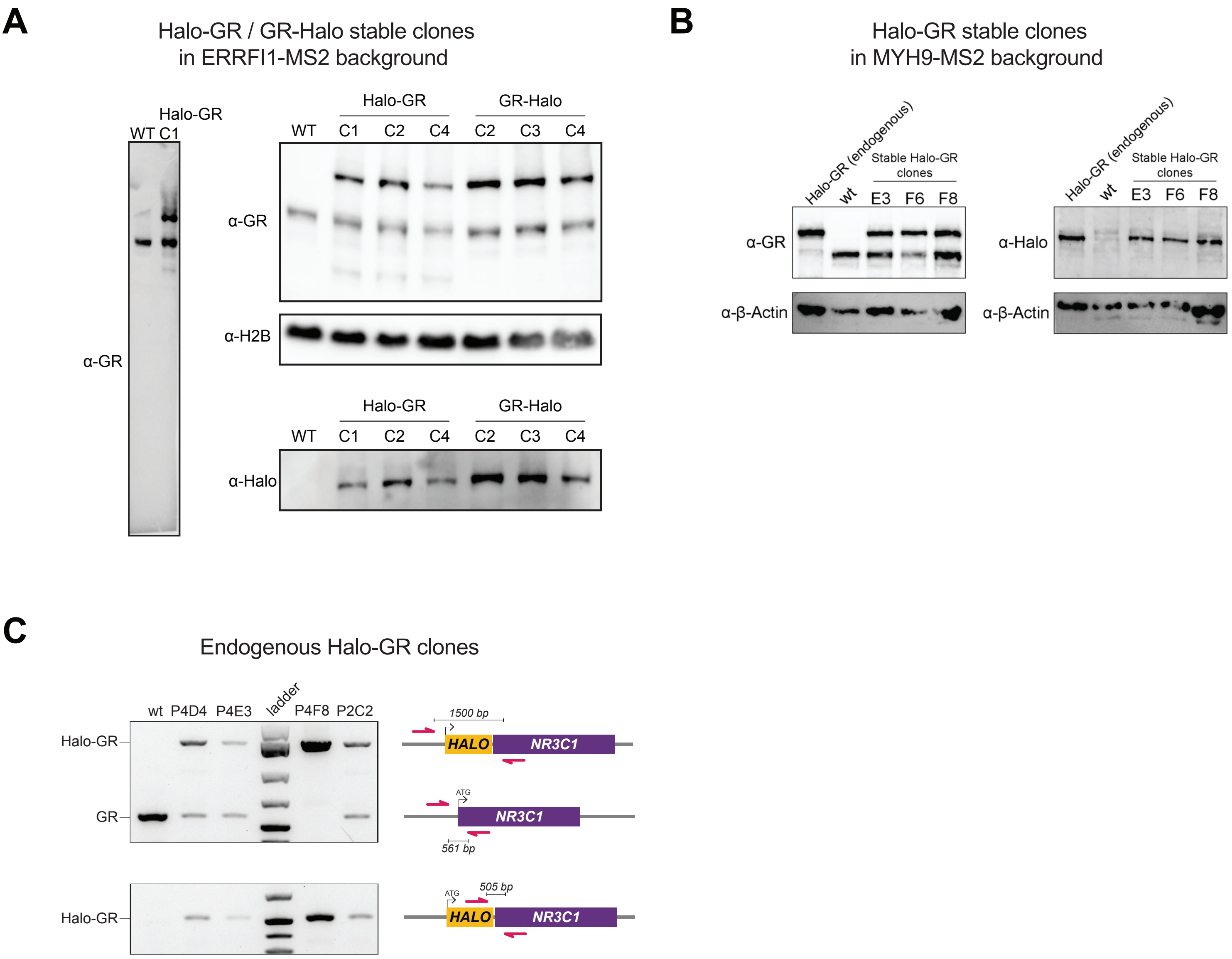


**Figure S2. Generation and validation of Halo-tagged GR cell lines.**

(A) Western blot validation of stable Halo-GR and GR-Halo clones generated by lentiviral integration in the ERRFI1-MS2 cell line. Blots were probed with antibodies against GR, H2B (loading control), and HaloTag. WT, parental untagged cells.

(B) Western blot validation of stable Halo-GR clones in the MYH9-MS2 cell line, probed with antibodies against GR, HaloTag, and β-Actin (loading control). An endogenous Halo-GR knock-in clone is included as a reference.

(C) Genotyping PCR validation of endogenous Halo-GR clones generated by CRISPR HDR knock-in of HaloTag at the N-terminus of *NR3C1*. Top: PCR primers flanking the insertion site detect both the Halo-GR fusion (upper band, ~1500 bp) and the untagged allele (lower band, ~561 bp). Bottom: a primer pair spanning the Halo-NR3C1 junction (~505 bp) confirms integration. WT, parental cells.

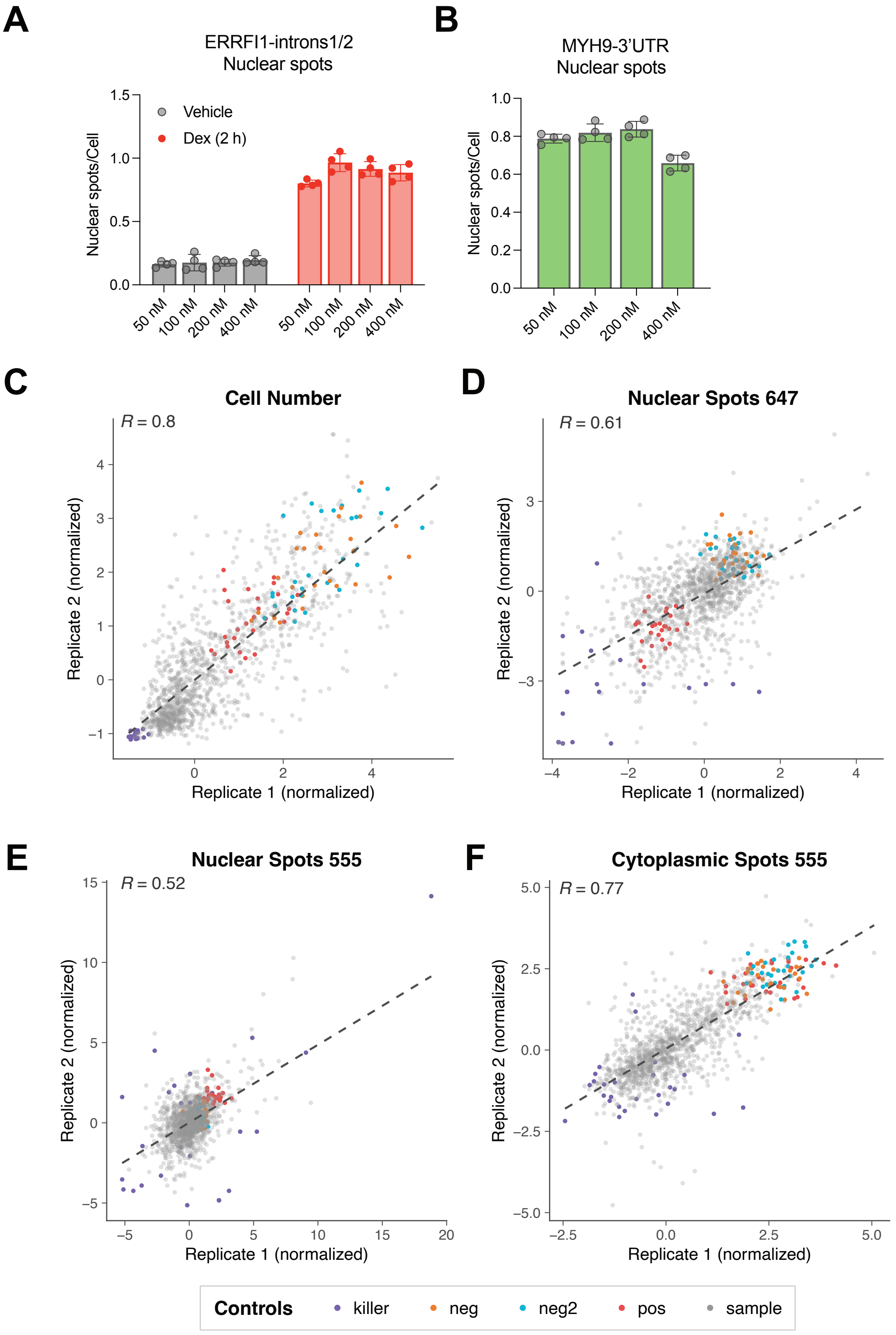


**Figure S3. Probe concentration optimization and replicate concordance of screen readouts.**
(A-B) Nuclear spots per cell for *ERRFI1* intron 1/2 (A) and *MYH9* 3'UTR probes at varying concentrations. (D-F) Scatter plots comparing normalized values between replicates for cell number (A), nuclear spots in the 647 channel (B), nuclear spots in the 555 channel (C), and cytoplasmic spots in the 555 channel (D). Dashed lines indicate linear regression fits with Pearson correlation coefficients (R) are shown for each readout.
